# Viral infection drives cell-intrinsic re-localization of the *C. elegans* immune-repressive STAT transcription factor STA-1

**DOI:** 10.64898/2026.06.24.734234

**Authors:** Lakshmi E. Batachari, Tyler D. Bechtel, Zhouxin Shen, Emily R. Troemel

**Affiliations:** School of Biological Sciences, University of California, San Diego, La Jolla, California, United States

**Keywords:** *C. elegans*, STAT transcription factor, nuclear localization dynamics, RIG-I-like receptor, viral infection, Orsay virus, intestinal immunity, innate immunity

## Abstract

Detection of viral infection leads to both cell-intrinsic and cell-extrinsic responses. In mammals, cell-intrinsic detection of viral infection leads to cell-extrinsic activation of STAT (Signal Transduction and Activators of Transcription) proteins, a family of transcription factors that promote anti-viral defense. In the nematode *C. elegans*, STA-1/STAT is a negative regulator of anti-viral defense, but it is not known if it acts cell-intrinsically or cell-extrinsically and whether it has functional domains conserved with mammalian STATs. Here we show that *C. elegans* STA-1 protein disappears from nuclei of cells infected with the natural viral pathogen, Orsay virus, but remains nuclear in uninfected cells, indicating a cell-intrinsic site of action. During viral infection, STA-1 forms cytoplasmic puncta that interact with the RNA viral sensor DRH-1, suggesting that DRH-1 helps restrain this immune-repressive factor. STA-1 overexpression causes increased susceptibility to viral infection, in a manner dependent on conserved residues important for DNA binding, nuclear localization and phosphorylation. Structural predictions indicate that STA-1 is most similar to STAT5 proteins in mammals, which have known immune-repressive roles. Our transcriptomic analysis demonstrates that *C. elegans* STA-1 regulates a general anti-pathogen program, including genes upregulated later during viral infection. Altogether, our findings provide insight into conserved and distinct features of STA-1 in *C. elegans*, indicating an ancient role for cell-intrinsic, immune-repressive STATs.

**Author Summary:** All living organisms must detect viral infections and mount a defense to survive. One major antiviral defense pathway in mammals is the interferon response, which involves sensing viral infection in one cell, and delivering an interferon ‘message’ to neighboring cells. These neighboring cells then turn on anti-viral defense gene expression using proteins called STAT transcription factors. We study anti-viral defense in the roundworm *C. elegans*, and in this study show that viral infected cells themselves use a STAT protein called STA-1, with perhaps a lesser role for STA-1 in neighboring cells, in contrast to mammals. We also extend on previous findings that STA-1 turns off anti-viral gene expression, and we analyze regions in the protein to demonstrate that STA-1 is bona fide transcription factor with an immune-repressive role. Structural prediction analysis of STA-1 indicates it is most similar to STAT5 in mammals, suggesting an ancient role for this protein as an immune-repressive factor acting directly in virally infected cells.

## Introduction

Detection of viral infection and activation of downstream anti-viral defense is critical for survival of all organisms. In vertebrates, RNA viruses in the cytosol are detected by RIG-I-like receptors (RLRs), which bind viral RNA ligands to activate cell-intrinsic transcription of type I and type III interferons[1, 2]. Interferons are secreted ligands, which bind to interferon receptors on neighboring, bystander cells. Cells that receive the interferon signal activate a signaling cascade through Signal Transduction and Activators of Transcription (STAT) proteins to upregulate transcription of hundreds of downstream interferon-stimulated genes (ISGs) that provide anti-viral defense[3]. The human genome has seven STAT transcription factors, with multiple, prominent roles in immunity[4]. While RLR signaling in infected cells followed by cell STAT activation in neighboring cells is the canonical model for mammals, there is evidence that STATs like STAT3 and STAT5 can be repressors of immunity. Furthermore, these immune-repressive STATs may have direct interactions with RLRs, although these roles are less well-understood.

The nematode *Caenorhabditis elegans,* like other invertebrates, lacks obvious interferon ligand/receptor homologs, but they have an anti-viral transcriptional response called the Intracellular Pathogen Response (IPR), with similarities to the type I/III interferon response[5]. The IPR is induced by infection with intracellular pathogens including the Orsay virus, a single-stranded, positive-sense RNA virus that is a natural intestinal pathogen of *C. elegans*. Like the type I/III interferon response, the IPR can be activated by a RLR homolog, called DRH-1 in *C. elegans*, which detects viral infection and then induces a downstream anti-viral transcriptional program[6]. During infection, DRH-1 protein transitions from being diffusely expressed in the cytoplasm to coalescing into discrete puncta, which colocalize with double-stranded RNA, a viral replication product known to activate vertebrate RLRs and the downstream type I/III interferon response[7]. These features are similar to mammalian RLRs, and suggest that DRH-1 puncta represent signaling complexes. Furthermore, structural analysis indicates DRH-1 binds dsRNA, based on analysis of the role of DRH-1 in anti-viral RNA interference[8], which has been shown to be distinct from the DRH-1 role in anti-viral transcriptional response[6].

How DRH-1 activates the IPR is not well-understood, as the *C. elegans* genome lacks obvious homologs of the signaling components that function downstream of mammalian RLR cell-intrinsic signaling, like MAVS and the transcription factors IRF-3/7 that upregulate transcription of interferon ligands[5]. *C. elegans* also lacks clear homologs for the signaling components that act downstream of the interferon receptor in mammals, such as the Janus kinases (JAKs) that activate STATs to induce transcription of ISGs. However, *C. elegans* does have two STAT homologs, STA-1 and STA-2[9]. Prior work demonstrated that STA-2 is important for inducing an anti-fungal transcriptional response in the epidermis[10], while STA-2 is not important for an antiviral response in the intestine[11]. Instead, STA-1 has been shown to be expressed in the intestine, and serves as a repressor for anti-viral immunity in this tissue[11]. Specifically, STA-1 binds a DNA region similar to the consensus binding sequence of human STAT proteins, and represses expression of viral response genes. Accordingly, *sta-1* mutants have increased resistance to viral infection. However, it is not known whether STA-1 acts in bystander uninfected cells, similar to canonical mammalian STAT signaling, or whether STA-1 acts in infected cells. Furthermore, it is not known whether the transcriptional activity of STA-1 is required for its immune-repressive function, nor is it known the stage of infection when STA-1 acts.

Here, we investigate the localization of STA-1 during infection and immune activation mediated by DRH-1, and probe the structure/function relationship of STA-1 protein domains in immunity. During viral infection, we observe the loss of STA-1 protein from the nuclei of virus-infected cells, but maintenance of STA-1 nuclear expression in bystander, uninfected cells. This behavior is distinct from canonical STAT signaling in mammals, where STAT signaling occurs in bystander cells. Furthermore, we demonstrate that STA-1 colocalizes with DRH-1 in the cytoplasm of virus-infected cells, consistent with our finding that STA-1 is physically associated with the signaling domain of DRH-1. Importantly, we show that STA-1 overexpression causes susceptibility to viral infection, and this immune-repressive capability of STA-1 is dependent on its nuclear localization, its DNA binding residues, and a conserved C-terminal tyrosine residue that is phosphorylated by the upstream activating JAK in mammals. Transcriptomic analysis reveals that *C. elegans* STA-1 represses a subset of genes induced during later stages of viral infection. Its anti-viral activity, however, appears to act upstream of viral replication, as we find that an Orsay viral replicon is not inhibited by *sta-1* RNAi, implicating STA-1 in a pre-replication step. Altogether, our work provides insight into STAT signaling in *C. elegans*, suggesting that the immune-repressive capability of STA-1 is cell-intrinsic, with parallels to STAT5 signaling in mammals.

## Results

### Viral infection causes loss of nuclear STA-1 expression in intestinal cells

In a previous study, we demonstrated that overexpression of the DRH-1 signaling domain, DRH-1(2CARD), was sufficient to induce Intracellular Pathogen Response (IPR) genes and confer antiviral resistance[7]. To determine factors that signal downstream of DRH-1, we performed co-immunoprecipitation/mass spectrometry (co-IP/MS) to identify proteins that interact with DRH-1(2CARD)::mScarlet (Fig. S1). Here, we found that STA-1 was significantly enriched in DRH-1(2CARD)::mScarlet protein fractions relative to an mScarlet control. Because previous work demonstrated that STA-1 acts as a transcriptional repressor of virus response genes, we investigated STA-1 protein behavior in the context of viral infection and DRH-1 signaling.

One hypothesis for STA-1 signaling is that it leaves the nucleus during infection, in order to de-repress immune gene expression. To investigate this hypothesis, we examined STA-1::GFP following infection with Orsay virus. Consistent with previous reports[11], we saw STA-1::GFP was enriched in the nuclei of intestinal cells in uninfected animals (Fig. 1A). In contrast, we saw a significant decrease in nuclear STA-1::GFP at 24 hours post-inoculation (hpi) in virus-infected cells, as indicated by FISH fluorescence (Figs. 1A, B). We also observed a significant decrease in the ratio of nuclear to cytoplasmic GFP fluorescence (N/C ratio) in virus-infected cells (Fig. 1C). Canonical STAT signaling in mammals occurs in a cell-extrinsic manner, whereby infected cells activate STAT in bystander uninfected cells. To assess whether the loss of nuclear localization occurred cell-extrinsically, we compared STA-1::GFP fluorescence in infected (FISH+) vs. uninfected (FISH-) cells within the same *C. elegans* animal. Notably, we found that STA-1::GFP was present in the nuclei of neighboring, uninfected cells (Fig. 1D), indicating that virus-triggered loss of nuclear STA-1 occurs in a cell-intrinsic manner.

**Fig. 1.**
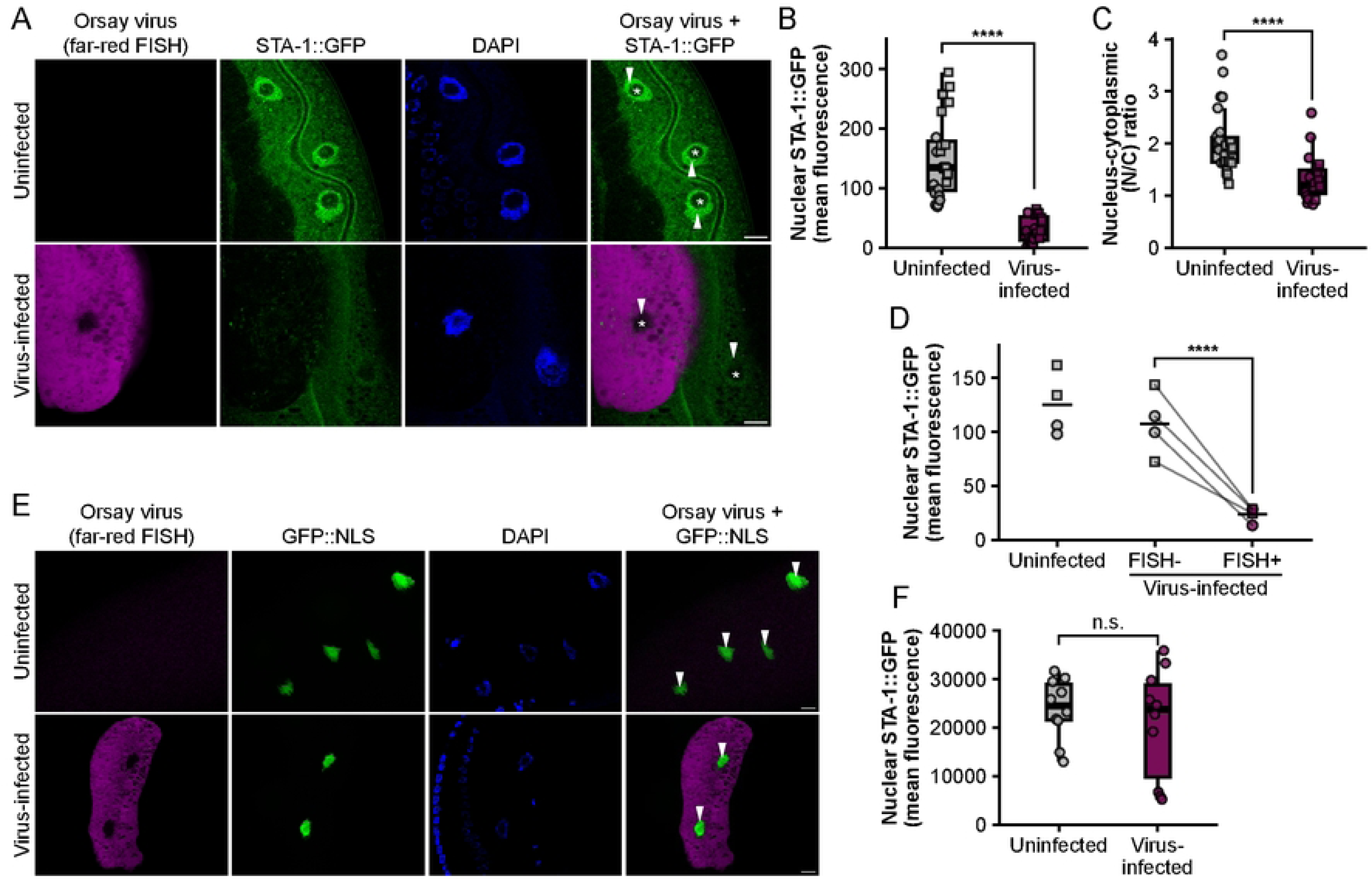
Viral infection causes loss of STA-1 protein in intestinal cell nuclei. (A) Representative images of uninfected or virus-infected adult animals at 24 hpi. Animals express endogenously tagged STA-1 (green) and were infected at the L4 stage. Viral infection was visualized using Quasar 670-conjugated (far-red) FISH probes targeting the Orsay virus genome. Nuclei were counterstained using DAPI (blue). White arrowheads indicate nuclei. Asterisks denote nucleoli. (B) Mean STA-1::GFP fluorescence intensity values in the nucleus of virus-infected (FISH+) vs. uninfected cells. (C) Ratio of nuclear-to-cytoplasmic STA-1::GFP fluorescence in virus-infected (FISH+) vs. uninfected cells. For (B) and (C), each dot represents one nucleus. Two independent experimental replicates were performed. For each replicate, 6-8 animals/treatment were analyzed with 1-3 nuclei quantified per animal. Different experimental replicates are represented by different dot symbols. (D) Mean STA-1::GFP fluorescence intensity in the nuclei of uninfected worms and virus-infected worms, containing both FISH+ and FISH-cells. Each dot represents one nucleus; data reflect two independent experimental replicates. Line segments connect distinct nuclei in the same worm. Horizontal crossbar represents the mean. A paired t-test was used to calculate p-values; ****p < 0.0001 (E) Representative images of viral infection in animals expressing GFP fused to a nuclear localization sequence (NLS). Viral genome was stained using Quasar 670-conjugated (far-red) FISH probes and nuclei were counterstained with DAPI (blue). (F) Quantification of mean GFP::NLS fluorescence in the nucleus of virus-infected vs. uninfected cells in (E). In (A) and (E), white arrowheads indicate nuclei and asterisks denote nucleoli. For (B), (C), and (F), horizontal lines in box and whisker plots represent median values, and the box reflects the 25th to 75th percentiles. A Mann-Whitney U test was used to calculate p-values; ****p < 0.0001. Scale bar = 10 µm for all images.

To determine the kinetics of STA-1::GFP localization, we examined animals at 12, 24, and 48 hpi. STA-1::GFP remained enriched in the nucleus at 12 hpi, whereas this enrichment was no longer present at 24 hpi and 48 hpi (Fig. S2). As a negative control, we assessed how viral infection impacted constitutively expressed GFP fused to a nuclear localization sequence (GFP::NLS). In virus-infected cells, there was no change in nuclear GFP expression compared to uninfected cells (Figs. 1E, F). These findings indicate that the loss of nuclear STA-1::GFP in infected cells is not a non-specific consequence of viral infection causing a general reduction in nuclear GFP expression.

Next, we used a viral replicon system to investigate whether viral replication alone was sufficient to cause the loss of nuclear STA-1::GFP. Here we used a previously established heat shock-inducible system to express the Orsay virus RNA1 genome segment, which contains a gene that encodes the RNA-dependent RNA polymerase (RDRP). Heat shock induction of wild-type RNA1, as part of the *virEx23* transgene, catalyzes RNA1 replication and induces the expression of viral response genes similar to those induced by native viral infection. At 24 h post-heat shock induction of RNA1, animals showed a trend towards reduced nuclear STA-1::GFP relative to non-induced controls (Fig. 2A, 2B). Heat shock alone resulted in increased, rather than decreased, nuclear STA-1::GFP (Fig. 2C, 2D). This finding suggests that the trend towards decreased STA-1::GFP is due to the RNA1 replicon and not heat shock itself. Both heat shock alone and heat shock induction of RNA1 resulted in loss of nuclear enrichment of STA-1::GFP, which is reflected by a decrease in the N/C ratio (Fig. 2E, 2F). In summary, expression of the Orsay virus RNA1 replicon drives a moderate reduction in nuclear STA-1::GFP. These findings suggest that the loss of nuclear STA-1::GFP during viral infection is, in part, due to replication of the viral genome.

**Fig. 2.**
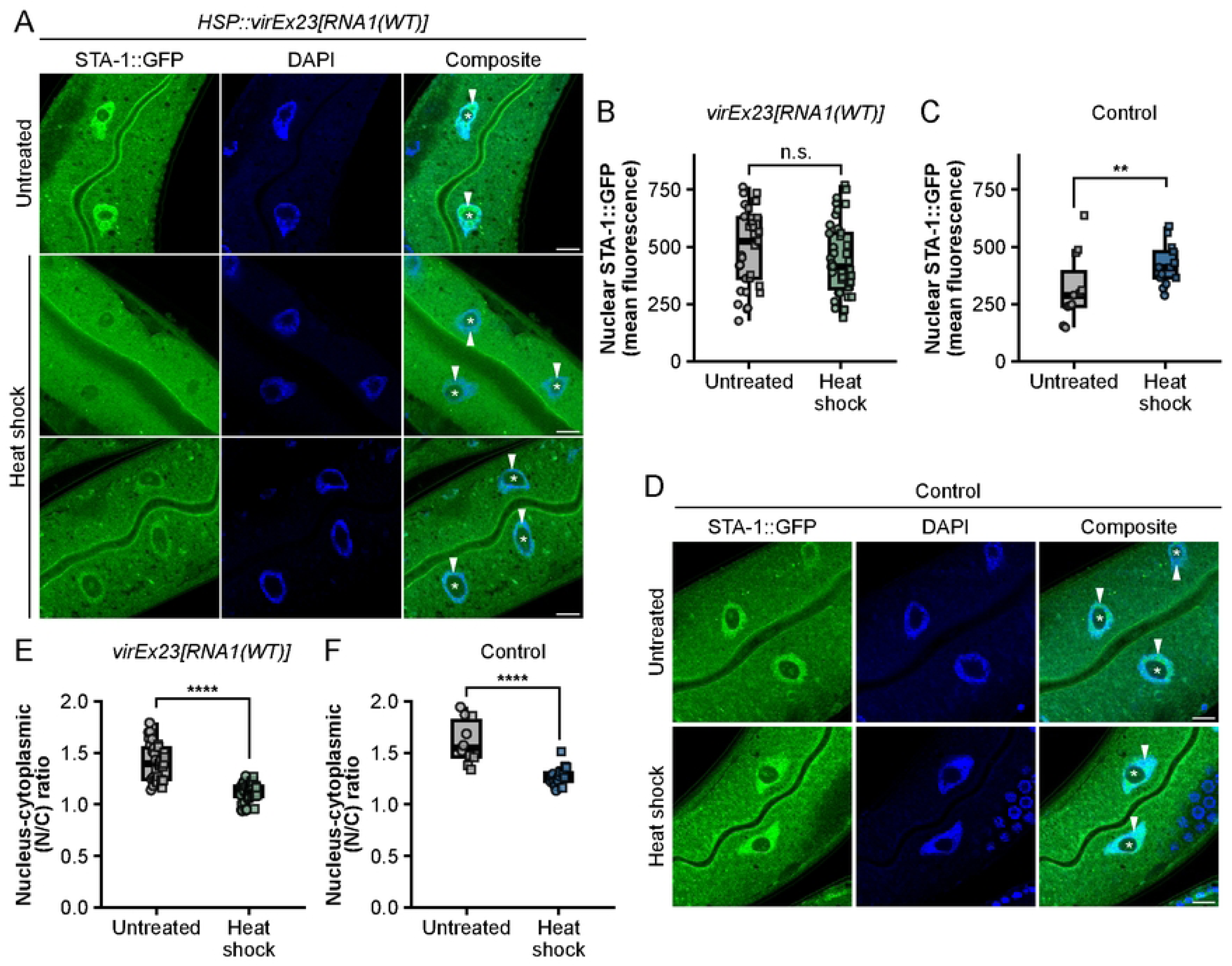
Orsay virus RNA1 replicon is sufficient to drive decreased nuclear enrichment of STA-1. (A) Representative images of endogenous STA-1::GFP in animals that express the *virEx23[HSP::RNA1]* transgene, which allows for heat shock-inducible expression of the Orsay RNA1 replicon. (B) Quantification of nuclear STA-1::GFP fluorescence in *virEx23[HSP::RNA1]* animals. (C) Quantification of nuclear STA-1::GFP fluorescence in control animals. (D) Representative images of endogenous STA-1::GFP in control animals. (E) Nuclear-cytoplasmic GFP fluorescence ratio in *virEx23[HSP::RNA1]* animals. (F) Nuclear-cytoplasmic GFP fluorescence ratio in control animals. For (B, C, E, F), each dot represents one nucleus. For each condition, data reflect two biological replicates (two plates) from one experimental replicate. Horizontal lines in box and whisker plots represent median values, and the box reflects the 25th to 75th percentiles. A Mann-Whitney U test was used to calculate p-values; ****p < 0.0001. White arrowheads indicate nuclei and asterisks denote nucleoli. Scale bar = 10 µm for all images.

### RIG-I-like receptor DRH-1 colocalizes with STA-1::GFP and drives loss of nuclear enrichment of STA-1::GFP

In addition to the loss of STA-1 from the nucleus, we found that a subset of virus-infected cells showed the formation of cytoplasmic, endogenous STA-1::GFP puncta (Fig 3A). In previous work, we demonstrated that the RIG-I-like receptor homolog DRH-1, which is a cytosolic sensor for Orsay virus, forms puncta in virus-infected cells [7]. Therefore, we asked whether the formation of STA-1::GFP and mScarlet::DRH-1 puncta occurred simultaneously during infection. To address this question, viral infections were performed in animals that express endogenously tagged STA-1::GFP and a single-copy mScarlet::DRH-1 transgene. Here, we found partial overlap between mScarlet::DRH-1 signal and STA-1::GFP signal (Fig. 3A), suggesting that DRH-1 and STA-1 may be recruited to some of the same cytoplasmic structures during infection.

**Fig. 3.**
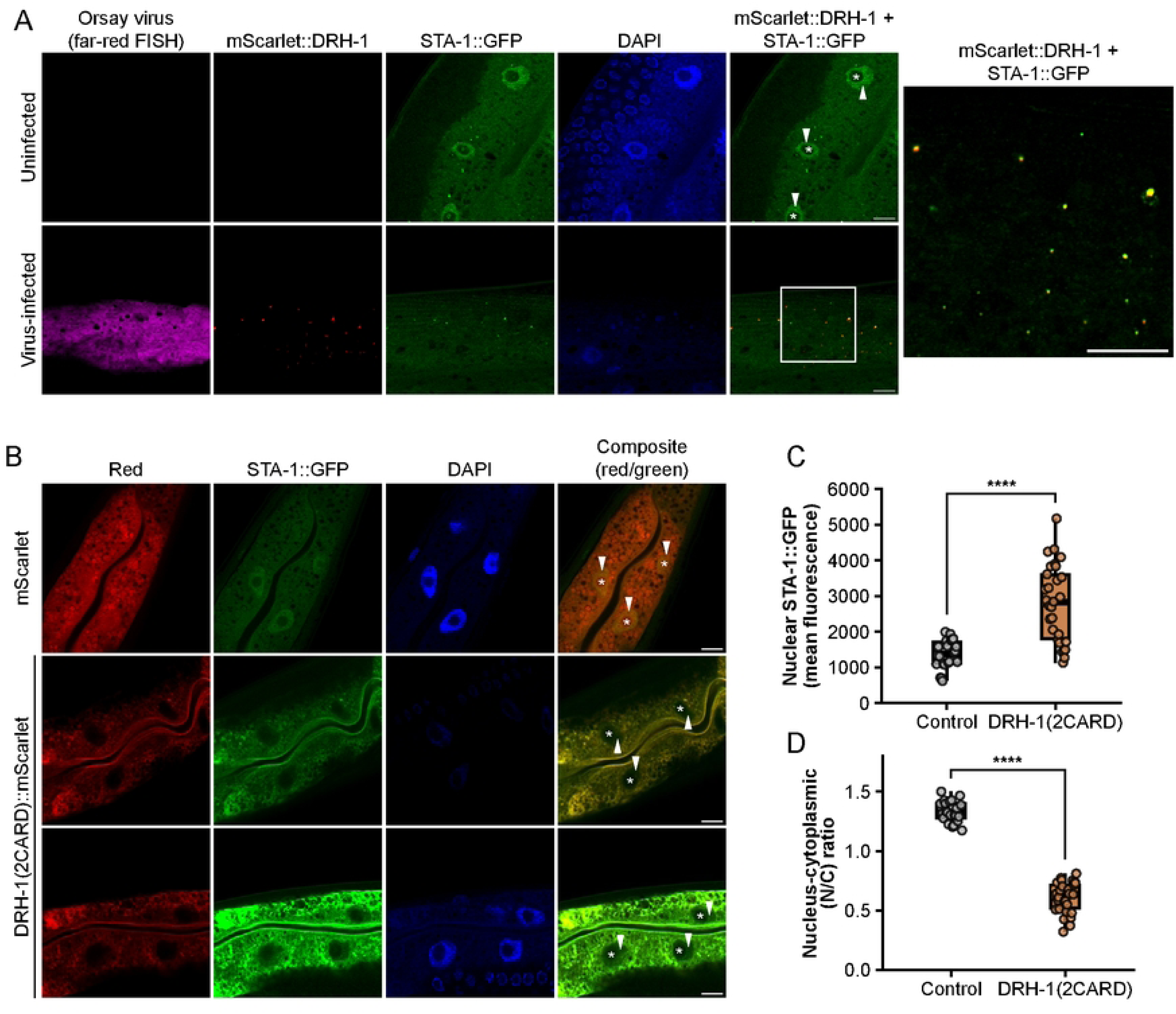
RIG-I-like receptor DRH-1 colocalizes with STA-1, and DRH-1(2CARD) signaling drives loss of STA-1 nuclear enrichment. (A) Representative images of uninfected or virus-infected animals with endogenously tagged STA-1::GFP (green) and an integrated single-copy mScarlet-tagged DRH-1 transgene (red). Animals were infected at the L4 stage for 24 h. Viral infection was visualized using Quasar 670-conjugated (far-red) FISH probes targeting the Orsay virus genome. Nuclei were counterstained with DAPI (blue). (B) Representative images of animals expressing endogenously tagged STA-1::GFP and DRH-1(2CARD)::mScarlet as an integrated, multi-copy transgene. (C) Mean STA-1::GFP fluorescence intensity values of nuclei in mScarlet vs. DRH-1(2CARD) animals. (D) Ratio of nuclear-to-cytoplasmic STA-1::GFP fluorescence in mScarlet vs. DRH-1(2CARD) animals. For (A) and (B), white arrowheads indicate nuclei and asterisks denote nucleoli. For (C) and (D), each dot represents one nucleus. 10 animals per strain were analyzed; 1-3 nuclei were scored per animal. Horizontal lines in box and whisker plots represent median values, and the box reflects the 25th to 75th percentiles. A Mann-Whitney U test was used to calculate p-values; ****p < 0.0001. Scale bar = 10 µm for all images.

Upon viral infection and induction of the RNA1 replicon, DRH-1 is required for the transcription of virus response genes, including genes that constitute the Intracellular Pathogen Response (IPR). Previously, we demonstrated that, similar to RLR signaling in mammals, DRH-1 contains an N-terminal tandem caspase activation and recruitment domain (2CARD) that is sufficient to induce the expression of IPR genes and confer anti-viral resistance[7]. Given that the loss of *sta-1* confers anti-viral resistance, we sought to determine whether DRH-1(2CARD)-mediated signaling alone was sufficient to trigger the loss of nuclear STA-1::GFP. Here, we generated a strain containing an integrated multi-copy transgene, in which the intestine-specific promoter *vha-6p* drives constitutive expression of mScarlet-tagged DRH-1(2CARD). This transgene was introduced into a strain containing endogenously tagged STA-1::GFP. To control for effects of mScarlet alone, we also generated animals that express intestine-specific mScarlet as part of an integrated multi-copy transgene. Upon DRH-1(2CARD)::mScarlet overexpression, we observed significantly elevated levels of nuclear STA-1::GFP (Fig. 3B, 3C). While total levels of STA-1::GFP are increased, we found that DRH-1(2CARD) animals exhibited a significant decrease in the N/C ratio (Fig. 3D). These findings suggest that DRH-1 signaling promotes STA-1 re-localization in the absence of infection.

### Immune-repressive role of STA-1 is dependent on nuclear localization and amino acids predicted to control DNA binding

Our results described above indicate that viral triggers, as well as immune signaling, cause a decrease in STA-1 nuclear enrichment. Thus, we hypothesized that the immune-repressive activity of STA-1 requires its nuclear enrichment and canonical transcription factor functions. Phylogenetic analysis of 229 full-length STAT sequences reveals that *C. elegans* STA-1 and STA-2 cluster separately from STATs in other organisms (Fig. S3). When we performed Foldseek analysis, however, we found that structural predictions of STA-1 indicated that it shares structural similarity with human STAT5a and STAT5b (Fig. S4). STAT5 has previously been shown to have immune-repressive functions[12] [13], and to bind RIG-I in a cell-intrinsic manner[14].

Based on the predicted Alphafold structure, STA-1 has domains characteristic of mammalian STAT proteins. STA-1 appears to contain a coiled-coil domain, a DNA binding domain, and a linker domain reminiscent of those found in STAT5a and STAT5b (Figs. 4A, S4). In order to perform structure/function analysis of STA-1, we cloned a construct containing wild-type STA-1, under the control of its native 5’ regulatory sequence. This wild-type STA-1::GFP construct was expressed from a multi-copy array in *sta-1(ok587)* loss-of-function mutants (Fig. 4B). Similar to what we observed with endogenously tagged STA-1, multi-copy STA-1(WT) protein levels were markedly decreased in virus-infected (FISH+) intestinal cells (Fig. 4B). We then assessed the ability of STA-1(WT) to rescue susceptibility to infection in *sta-1(ok587)* mutants, which are highly resistant to infection. Quantification of infection rates by FISH revealed that STA-1(WT) overexpression in two independent transgenic lines (*jyEx452, jyEx453*) was sufficient to rescue susceptibility to viral infection (Figs. 4D, E). Of note, STA-1(WT) animals displayed increased susceptibility relative to wild-type N2 animals (Fig. 4D).

**Fig. 4.**
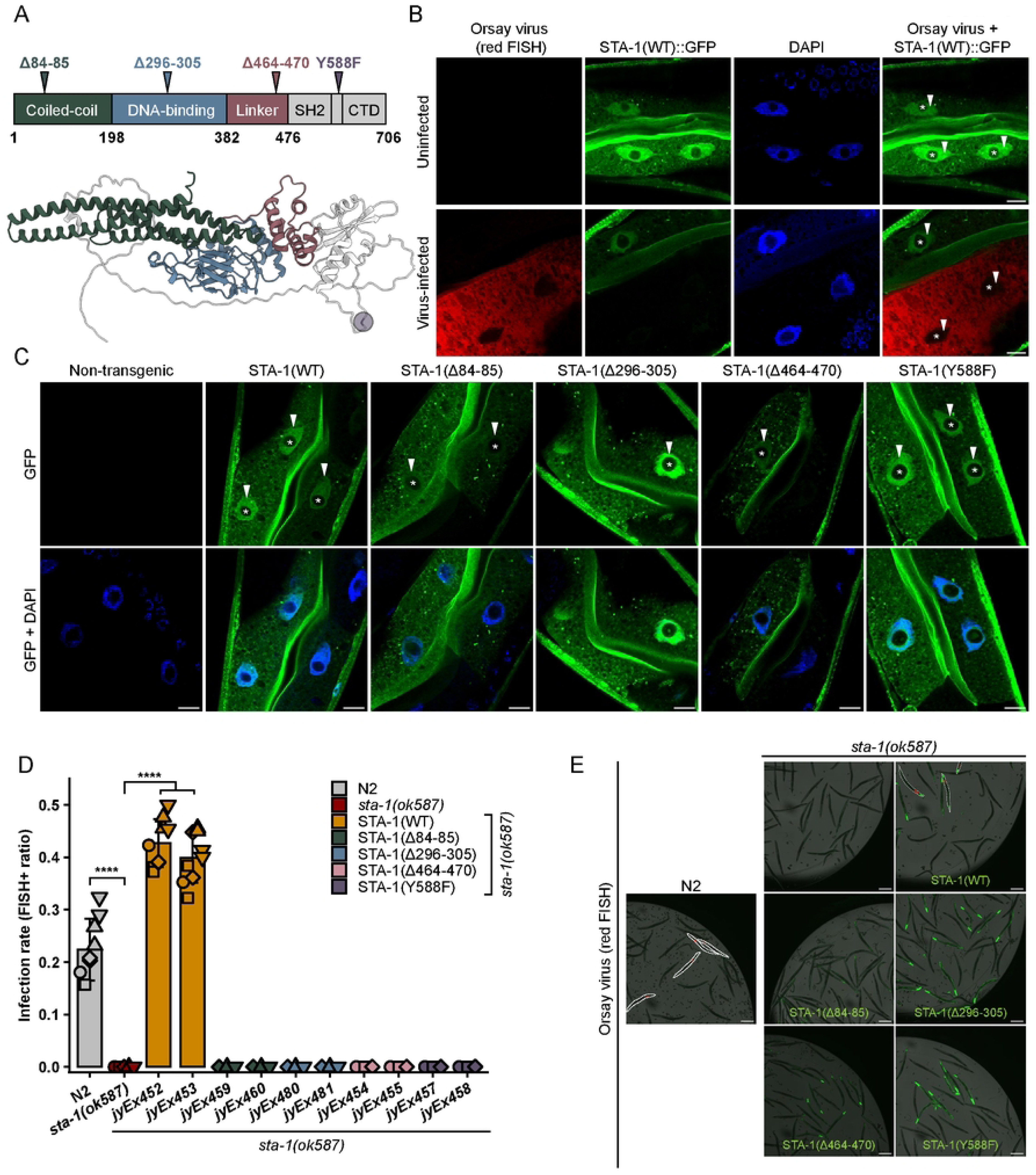
Immune-repressive activity of STA-1 is dependent on canonical STAT transcription factor functional domains. (A) Alphafold prediction of STA-1 protein structure showing the following putative domains: coiled-coil (green), DNA binding (blue), and linker (pink). The C-terminal tyrosine residue is highlighted in purple. (B) Representative images of uninfected or virus-infected animals that express STA-1::GFP from a multi-copy, extrachromosomal array. Viral infection was visualized using CF610-conjugated (red) FISH probes targeting the Orsay virus genome. Nuclei were counterstained with DAPI (blue). (C) Representative images of *sta-1(ok587)* animals that have either no transgene or a multi-copy, extrachromosomal transgene that encodes for GFP-tagged WT or mutant STA-1 protein. Nuclei were counterstained with DAPI (blue). For (B) and (C), white arrowheads indicate nuclei and asterisks denote nucleoli. Scale bar = 10 µm. (D) Viral infection rates for wild-type N2, *sta-1(ok587)* mutants, and various STA-1 extrachromosomal transgenic lines. Two extrachromosomal lines each were tested for STA-1(WT) and mutant constructs. For wild-type N2, *sta-1(ok587),* and STA-1(WT) lines (*jyEx452* and *jyEx453*), nine biological replicates (n = 9 plates) were scored over five independent experimental replicates. For all other lines, five biological replicates (n = 5 plates) were scored over three independent experimental replicates. Each dot represents a biological replicate of a minimum of 200 worms scored. Different experimental replicates are represented by different dot symbols. A pairwise *t-*test was used to determine significant differences relative to *sta-1(ok587)*; ****p < 0.0001. Mean values are represented by bar height; error bars represent the standard deviation. (E) Representative images of animals quantified in (D). Viral infection was visualized using CF610-conjugated (red) FISH probes targeting the Orsay virus genome. Infected animals are outlined by dotted white lines. Scale bar = 200 µm.

To determine whether nuclear localization of STA-1 is required for its immune-repressive effects, we deleted two amino acids (aa) in a conserved region found in the STA-1 coiled-coil domain (Δ84-85); the homologous region in human STAT3 functions as a nuclear localization sequence (NLS)[15]. When expressed in a *sta-1(ok587)* background, STA-1(Δ84-85) showed loss of nuclear enrichment in intestinal cells (Fig. 4C), indicating that this region is also required for nuclear localization in *C. elegans*. The STA-1(Δ84-85) mutant was unable to rescue susceptibility to virus, suggesting the nuclear localization is required for the immune-repressive function of STA-1 (Figs. 4D, E).

We next examined residues predicted to be involved in DNA binding. We generated a 10 aa deletion mutation in the putative DNA binding domain (Δ296-305), in order to remove conserved lysine residues that in human STAT1 are required for binding core STAT target sequences[16]. Here, the localization pattern of STA-1(Δ296-305) phenocopied STA-1(WT), showing clear nuclear localization, indicating these residues are not important for nuclear localization of STA-1 (Fig. 4C). This finding is in contrast to human STAT1, where these residues are required[16]. A second deletion (Δ464-470) in another STAT1-homologous region[17] reduced both nuclear enrichment and overall protein expression (Fig. 4C). Both mutants failed to rescue viral susceptibility (Figs. 4D, E). These results indicate that the immune-repressive function of STA-1 requires not only nuclear localization, but also DNA-binding activity.

A highly conserved feature of STAT transcription factors is a tyrosine residue found in the C-terminus[4]. In classical STAT signaling, phosphorylation at this tyrosine residue mediates STAT dimerization and translocation into the nucleus. To test the role of this conserved tyrosine in *C. elegans* STA-1, we generated a tyrosine to phenylalanine point mutation (Y588F) at the C-terminal tyrosine residue, which is present in all human STATs. Unexpectedly, STA-1(Y588F) retained its nuclear enrichment, with localization patterns that resembled the wild-type protein (Fig. 4C). In contrast to STA-1(WT), however, STA-1(Y588F) was unable to rescue viral infection rate to wild-type N2 levels (Figs. 4D, E). Taken together, these mutant phenotypes suggest that the immune-repressive function of STA-1 is dependent on canonical STAT transcription factor signaling elements, including nuclear localization, DNA-binding activity, and a conserved C-terminal tyrosine.

### RNAi against *sta-1* fails to restrict a viral replicon, but upregulates genes induced by viral replication and DRH-1-mediated signaling

Given the robust viral restriction of *sta-1* mutants shown above, we next examined whether this restriction occurs pre- or post-viral replication. To address this question, we performed RNA-seq following RNAi against *sta-1* in a *virEx23[RNA1(WT)]* heat shock-inducible viral replicon background (using the system shown in Fig. 2). As a control, we compared these animals to those carrying *virEx26[RNA1(mt)],* which harbors a catalytic mutation abolishing RDRP polymerase activity, and thus RNA1 replication. We asked whether *sta-1* RNAi inhibits viral replication by quantifying RNA-seq reads mapped to the Orsay virus genome. RNA1 transcript levels were significantly higher in *virEx23[RNA1(WT)]* animals relative to *virEx26[RNA1(mt)]* animals, validating the system (Fig. 5A). Notably, we found that RNA1 transcript levels were slightly higher in animals treated with *sta-1* RNAi compared to control. Therefore, STA-1 likely regulates a pre-replication step, as *sta-1* defective animals do not appear to restrict replication of the viral genome.

**Fig. 5.**
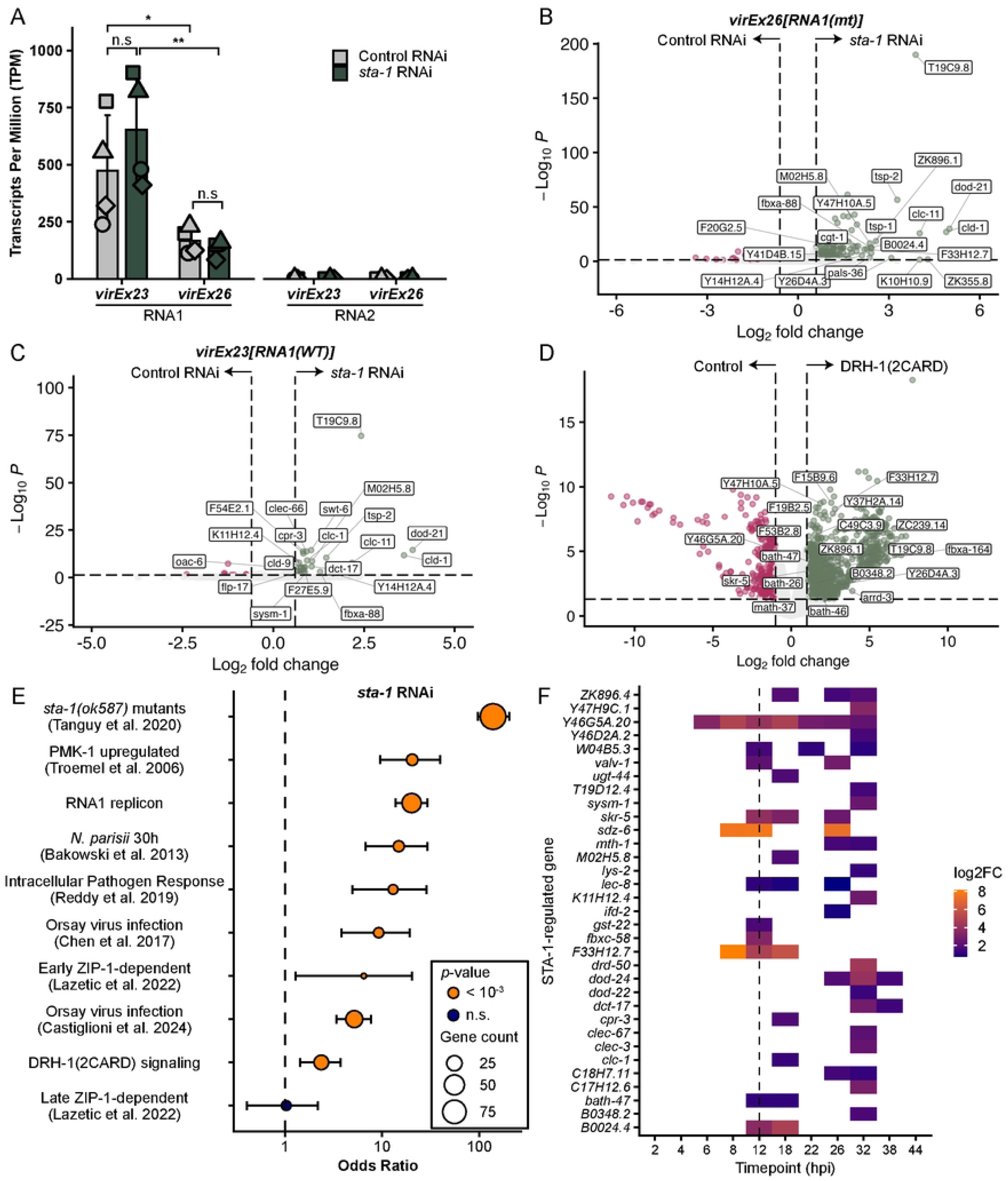
STA-1 negatively regulates genes that are induced by viral replication and DRH-1(2CARD)-mediated signaling. (A) Transcripts per Million (TPM) for Orsay viral RNA1 and RNA2 in RNA-seq samples. Each dot represents one experimental replicate, with different dot shapes corresponding to an experimental replicate. *virEx23[RNA1(WT)]* and *virEx26[RNA1(mt)]* animals were treated with either *sta-1* RNAi (green) or control RNAi (gray). Bars represent mean values; error bars indicate standard deviation. A Mann-Whitney U test was used to calculate p-values; *p < 0.05, **p < 0.01. (B) Volcano plot of differentially expressed genes (DEGs) in animals treated with *sta-1* RNAi vs. control RNAi in the absence of viral replication (*virEx26[RNA1(mt)]*). (C) Volcano plot of DEGs in animals treated with *sta-1* RNAi vs. control RNAi in the presence of viral RNA1 replication (*virEx23[RNA1(WT)]*). Upregulated (green) and downregulated (magenta) genes determined by |log2FC| > 0.6 and p < 0.05. For (B) and (C), top 20 upregulated genes are labeled. (D) Volcano plot of DEGs in DRH-1(2CARD) overexpressing animals. Upregulated (green) and downregulated (magenta) genes determined by |log2FC| > 1 and p < 0.05. Labels denote genes that are also upregulated by *sta-1* RNAi. (E) Odds ratio from Fisher’s exact test comparing upregulated genes from *sta-1* RNAi treatment with various gene sets listed on the y-axis. Error bars indicate 95% confidence interval. Significant associations (p < 0.001) are indicated by orange dots. Dot size reflects overlapping gene count. (F) Heatmap showing genes upregulated by *sta-1* RNAi (this study) that are also upregulated during infection time course (RNA-seq data from Castiglioni et al., 2024). Fold change values calculated relative to uninfected control.

We next investigated how STA-1 regulates the transcriptional response to viral replication. RNA-seq profiling previously revealed that *sta-1(ok587)* mutants have upregulation of Orsay virus response genes in the uninfected state[11]. However, the highly resistant phenotype of *sta-1(ok587)* mutants poses a challenge to assess the transcriptional response to viral infection, because *sta-1* mutants have a much lower ‘dose’ of virus compared to wild-type animals. To circumvent this issue, we analyzed the impact of *sta-1* RNAi with RNA-seq analysis of the viral replicon system. Differential expression analysis at 24 h post heat-shock revealed that, in the absence of RNA1 replication (*virEx26[RNA1(mt)]*), knockdown of *sta-1* resulted in the upregulation of 153 differentially expressed genes (DEGs) relative to treatment with control RNAi (Fig. 5B). This transcriptional de-repression is consistent with published RNA-seq data from *sta-1(ok587)* mutants[11]. In the context of the viral replicon (*virEx23[RNA1(WT)]*), we observed a marked reduction in number of upregulated genes (35 total) following *sta-1* RNAi (Fig. 5C). Given that *sta-1* RNAi treated animals experience similar or even greater levels of RNA1, this decrease in the number of upregulated genes provides support that the transcriptional response to viral replication is directly regulated by *sta-1*.

Genes upregulated by *sta-1* RNAi (in the absence of viral replication) were enriched for genes belonging to the functional category ‘stress response:pathogen’, as analyzed by WormCat[18] (Fig. S5A). We also profiled genes in the context of viral replication without *sta-1* RNAi treatment by comparing *virEx23[RNA1(WT)]* to *virEx26[RNA1(mt)]*, where both groups were treated with control RNAi (Fig S5B). Here, we identified 447 upregulated genes, a subset of which were also classified under the ‘stress response:pathogen’ functional category (Fig. S5C). Notably, we identified 45 genes that are significantly upregulated by both *sta-1* RNAi and RNA1 replication (Fig. S5D). Seven of these overlapping genes are characterized as IPR genes: *pals-36, Y47H10A.5, F15B9.6, skr-5, Y46G5A.20, Y37H2A.14,* and *dod-23.* Next, to investigate the relationship between STA-1 and DRH-1(2CARD), we performed RNA-seq comparisons of DRH-1(2CARD)-expressing animals compared to control (Fig. 5D). Here we saw upregulation of 1335 genes, 22 of which were also upregulated upon *sta-1* RNAi (Fig. S5E).

To assess the broad relevance of STA-1-regulated genes in immunity, we compared genes upregulated by *sta-1* RNAi with genes upregulated in the context of other pathogen and immune responses (Fig. 5E). The 22 genes induced by both *sta-1* RNAi and DRH-1(2CARD) exhibited significant overlap (Fig. 5F). As measured by the number of overlapping genes, the largest overlap occurred with the upregulated gene set reported for *sta-1(ok587)* mutants, followed by the upregulated gene set for RNA1-induced genes and Orsay virus infection-induced genes (Castiglioni et al., 2024), respectively. We also observed modest, yet significant, overlaps between STA-1 regulated genes and other transcriptional defense programs, including PMK-1/p38 pathway genes and canonical IPR genes. During viral infection, activation of a subset of IPR genes is dependent on the transcription factor ZIP-1, with some genes requiring ZIP-1 at an early induction phase while others require ZIP-1 at a later phase. Moreover, *zip-1* mutants are more susceptible to infection[19, 20]. We found that genes negatively regulated by STA-1 are largely independent of ZIP-1-regulated genes. In summary, these findings support a model in which STA-1 represses a more general pathogen defense program.

Our findings related to STA-1::GFP localization indicate that STA-1 is lost from the nucleus at an infection timepoint between 12 and 24 hpi (Fig. S2). To examine the temporal dynamics of STA-1-mediated gene regulation over the course of infection, we compared upregulated genes from RNA-seq analysis of *sta-1* RNAi-treated animals (this study) with upregulated genes from a published infection time course spanning 2 to 44 hpi[21]. A subset of STA-1-regulated genes exhibited dynamic induction patterns during infection, with the majority of STA-1-regulated gene expression occurring between 12 and 32 hpi (Fig. S5F). The proportion of virus response genes negatively regulated by STA-1 peaks at 18 hpi (Fig. S5F). This timing is consistent with our observation that STA-1 disappears from the nucleus at a timepoint after 12 hpi (Fig. S2). Taken together, these findings support a model in which the loss of nuclear STA-1 leads to transcriptional de-repression of infection response genes at later stages of infection.

## Discussion

Our findings indicate that *C. elegans* STAT homolog STA-1 functions in a cell-intrinsic manner to de-repress anti-viral gene transcription during viral infection (Fig. 6). STA-1 exhibits loss of nuclear localization in virus-infected cells, but remains nuclear in neighboring, uninfected cells. STA-1 colocalizes with viral sensor DRH-1 in the cytoplasm of infected cells, consistent with the physical interaction observed between DRH-1(2CARD) and STA-1 from co-IP/MS analysis.

**Fig. 6.**
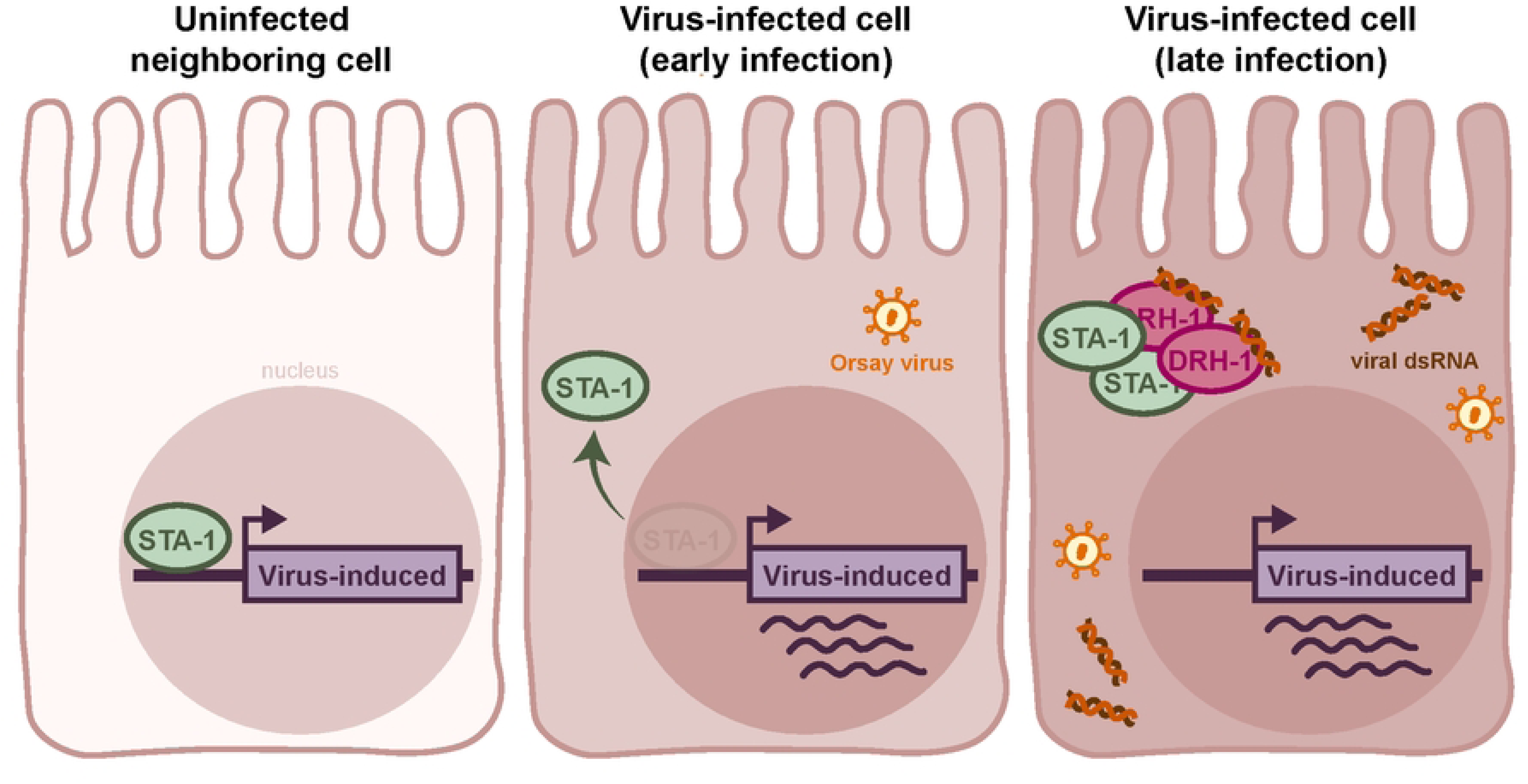
Model Figure.

Through structure/function studies, our results indicate that the anti-viral function of STA-1 is dependent on nuclear localization, conserved DNA binding residues, and a conserved C-terminal tyrosine residue. RNA-seq analyses using the Orsay virus replicon system indicate that STA-1 regulates a pre-replication stage of viral infection, but that the transcriptional de-repression of antiviral genes coincides with nuclear exclusion of STA-1 later during infection.

Contrary to the cell-extrinsic framework for mammalian STAT signaling, our data support a model in which a RIG-I-like receptor and STAT protein interact in a cell-intrinsic manner (Fig. 6). Our findings mirror a few recently described instances of RIG-I and STAT interacting in a cell-intrinsic pathway. For example, recent work describes ‘non-canonical’ modes of STAT signaling resulting from direct protein-protein interactions between RIG-I and STAT3[22]. In the absence of infection, RIG-I associates with STAT3 in T cells, thus preventing the phosphorylation and immune-repressive function of STAT3. In a model of colon cancer, a direct interaction between RIG-I and STAT3 attenuated tumor growth by inhibiting transcription of a STAT3 target gene[23]. Mechanistic studies in a murine tumor model implicate RIG-I and STAT5a in a cell-intrinsic pathway, whereby RIG-I represses STAT5a activity in CD8+ T cells[14]. Of note, the examples highlighted above describe interactions between RIG-I and STATs in a non-infection context. Findings from our study highlight a novel example of a cell-intrinsic RLR-STAT interaction occurring in the context of infection.

Another intriguing observation from this study is the requirement of the C-terminal tyrosine (Y588) for STA-1 function, but not for its nuclear localization. In canonical STAT signaling, JAK tyrosine kinases control phosphorylation of the conserved tyrosine residue to drive STAT dimerization and nuclear import[24]. Findings from cell culture have demonstrated that unphosphorylated mammalian STAT5 can be found in the nucleus[25]; of note, our Foldseek analysis indicated that *C. elegans* STA-1 is closest in predicted structure to STAT5. *C. elegans* lacks homologs to mammalian JAK tyrosine kinases and it is not clear which kinase performs analogous functions in phosphorylating STA-1. One candidate is the tyrosine kinase SID-3, as *sid-3* mutants have previously been shown to share similar phenotypes with *sta-1* mutants [11, 26]. Analysis of the interaction between SID-3 and STA-1 can be the subject of future studies.

What might drive the loss of STA-1 nuclear expression in virus-infected cells? Orsay virus infection disrupts intestinal cell morphology, including reorganization of intermediate filaments located at the apical membrane[27]. Our imaging indicates that STA-1 can be found at the apical membrane, which could be a key site for signaling. *C. elegans* has another STAT homolog, STA-2, that functions as a transcriptional activator of anti-microbial (AMP) genes in response to fungal infection[10] and epidermal wounding[28]. Under basal conditions, STA-2 associates with protein complexes, known as hemidesmosomes, located at the apical membrane of epidermal epithelial cells. Upon epidermal wounding, STA-2 dissociates from the hemidesmosomes and re-localizes to the nucleus, resulting in the transcription of target AMP genes[28]. Perhaps structural remodeling of the apical membrane in virus-infected intestinal cells retains STA-1 at this location, to prevent from shuttling back into the nucleus.

In this study, we describe a cell-intrinsic role for STA-1 that builds upon its initially documented role as a transcriptional repressor of anti-viral genes[11]. Findings in *C. elegans* contribute to a growing list of examples in which STAT proteins function as negative regulators of immune genes. In the context of mouse B cell development, DNA binding by STAT5 tetramers was associated with epigenetic repression of the target genes[12]. In CD4+ T helper cells, IL-2 signaling promoted STAT5 binding at the *Il17* locus to repress transcription and inhibit Th17 differentiation in a mouse model[13]. Given that *C. elegans* STA-1 is predicted to have the most structural similarity to STAT5 in mammals, our findings suggest an ancient role for STAT5-like transcription factors in the last common ancestor of mammals and *C. elegans*.

## Materials and Methods

### *C. elegans* Maintenance

*C. elegans* strains were maintained on Nematode Growth Media (NGM) agar plates containing streptomycin-resistant *Escherichia coli* OP50-1, unless otherwise specified. S1 Table lists all strains used in this study.

### Molecular cloning and transgenic *C. elegans* strains

For *sta-1* rescue experiments, the *sta-1* coding sequence was amplified from cDNA. Genomic DNA was used to amplify a 1432 bp sequence upstream of the *sta-1* start codon. Both the *sta-1* CDS and the 5’ upstream sequence were cloned into the plasmid pET848 to append an HA tag at the STA-1 C-terminus, followed by the *unc-54* 3’utr using Gibson assembly to generate pET850 [*sta-1p::sta-1::GFP::HA::unc-54 3’utr*] with the NEBuilder HiFi DNA Assembly kit (New England Biolabs). To clone STA-1 mutant constructs, mutations were introduced into *sta-1* cDNA of pET850 using the Q5 site-directed mutagenesis kit (New England Biolabs). All constructs were injected at 50 ng/μl along with 50 ng/μl filler DNA (pBluescript) into *sta-1* mutant animals to create extrachromosomal array transgenic lines.

For proteomics analysis, the *drh-1(2CARD)::mScarlet::3xFLAG* and *mScarlet::3xFLAG* transgenes were integrated using UV/psoralen. Briefly, pET770 and pET786 were injected into wild-type N2 worms at 100 ng/ul to generate extrachromosomal array lines. To integrate the transgene, L4 animals were incubated in 1 mg/mL of trimethyl psoralen for 15 min. at room temperature in the dark. Animals were plated onto unseeded NGM plates, UV-irradiated using a Stratalinker, and recovered overnight at 16°C. P0 animals were transferred onto fresh plates. F1s were cloned, followed by F2s cloned from plates with >75% transgenic worms. Integration was determined by identifying F2s that produced 100% transgene-containing progeny.

All plasmid sequences were validated by whole-plasmid sequencing. DNA constructs used in this study can be found in S2 Table. Primer sequences are listed in S3 Table.

### Synchronization of *C. elegans*

To synchronize development, gravid adult animals were washed off plates with M9 media and transferred into a 15 ml conical tube. Worms were pelleted at 3,000 rpm for 30 sec and resuspended in 3 ml of M9 and 1 ml of bleaching solution (500 μl of 5.65–6% sodium hypochlorite solution and 500 μl of 2 M NaOH). After most adults had dissolved, M9 was added to a final volume of 15 ml. Tubes were centrifuged and supernatant was discarded. Released embryos were washed an additional four times with 15 ml of M9 and resuspended in a final volume of 3 ml of M9. Embryos were incubated at 20 °C with continual rotation for 20-24 h to hatch L1s.

### Nuclear GFP fluorescence and N/C ratio measurements

Quantification of nuclear GFP was performed in QuPath. The wand tracing tool was used to outline the nucleus (excluding the nucleolus) based on DAPI fluorescence. The following wand settings were used: brush diameter = 8, point radius = 5, wand smoothing = 8, and wand sensitivity = 8. Mean GFP fluorescence was calculated by calculating intensity features using the following parameters: “pixelSizeMicrons” = 2 and “tileSizeMicrons” = 25.

To quantify cytoplasmic fluorescence, the cell detection tool was used to outline the cytoplasmic region bordering each nucleus such that the area of the cytoplasm is equivalent to that of the nucleus. The following parameters were used: “requestedPixelSizeMicrons”: 1.0, “backgroundRadiusMicrons”: 1000, “backgroundByReconstruction”: true, “medianRadiusMicrons”: 0, “sigmaMicrons”: 5, “minAreaMicrons”: 12, “maxAreaMicrons”: 1000, “threshold”: 23, “watershedPostProcess”: true, “cellExpansionMicrons”: 3, “includeNuclei”: true, “smoothBoundaries”: true, “makeMeasurements”: true.

### Orsay virus infections

All infections were performed on synchronized animals at the L4 stage, exposed to a mixture of OP50-1 bacteria, M9, and virus filtrate for 24 h at 20 °C, unless otherwise specified. All infections used the same batch of virus filtrate, prepared as previously described[29].

To visualize viral infection, animals were washed in M9 and fixed in 4% paraformaldehyde for 15 min. Following fixation, worms were incubated at 47 °C overnight with CF610- or Quasar 670-conjugated FISH probes that target Orsay virus RNA1 and RNA2 (Biosearch Technologies). Infection rate was assessed by the presence of FISH fluorescent signal. A minimum of two biological replicates (two plates, each containing 1000 animals) were performed per genotype per experiment.

### RNAi

RNAi was performed via the feeding method. RNAi bacterial clones (*sta-1* and L4440 empty vector control) were inoculated in 5 ml LB containing 50 μg/ml carbenicillin and incubated in a 30 °C shaking incubator (250 rpm) for 19.5 h. Overnight cultures were seeded onto NGM plates supplemented with 5 mM IPTG and 1 mM carbenicillin. Seeded RNAi plates were incubated at room temperature for 4 days.

### Heat shock induction of *HSP::*RNA1

Worms carrying the extrachromosomal arrays *virEx23[HSP::RNA1(wt), myo-2p::YFP]* or *virEx26[HSP::RNA1(mt), myo-2p::YFP]* were synchronized by bleaching. Hatched L1s were sorted based on expression of co-injection marker *myo-2p*::YFP using a COPAS Biosort instrument (Union Biometrica). For each treatment, 2000 YFP+ L1s were plated onto RNAi plates (two plates, each containing 1000 L1s). Worms were grown on RNAi plates at 20°C for 46 h until animals reached the L4 stage. To induce expression of RNA1(wt) from *virEx23* and RNA1(mt) from *virEx26,* synchronized L4s were exposed to a 2 h heat shock at 34°C, followed by a 24 h recovery at 20°C.

### RNA-seq sample preparation

For *virEx23[RNA1(WT)]* and *virEx26[RNA1(mt)]* samples, synchronized adult worms were collected at 24 h post-heat shock (see above). For DRH-1(2CARD) samples, *jyIs37; rde-1(ne300)* and *jyIs41; rde-1(ne300)* animals were synchronized by bleaching; 2000 L1s were plated onto seeded 10 cm plates and grown for 72 hours.

Prior to RNA isolation, animals were washed three times in M9 buffer. RNA was isolated using TRI Reagent (Molecular Research Center, Inc.) and 1-bromo-3-chloropropane (BCP) (Molecular Research Center, Inc.), followed by isopropanol and ethanol washes. Isolated RNA was resuspended in nuclease free water. RNA was further purified using the RNeasy cleanup kit with on-column DNase I digestion (Qiagen).

### RNA-seq

Generation of cDNA libraries and paired-end sequencing were conducted by the IGM Genomics Center, University of California, San Diego, La Jolla, CA. Sequencing was performed using the Illumina NovaSeq X Plus.

### RNA-seq data analysis

Alignment of RNA sequencing reads was performed using STAR version 2.7.11b[30] with the parameters: --runThreadN 24 --readFilesCommand zcat --outSAMtype BAM SortedByCoordinate --outFileNamePrefix. Reads were aligned to the WormBase WS279 reference genome. Read counts were generated using featureCounts with parameters: -p -s 2 -a. Normalization of sequencing reads and differential gene expression analysis were performed using the DESeq2 package in R[31]. For sequencing of *virEx23/virEx26,* differentially expressed genes were filtered out based on adjusted *P* < 0.05 and |log2 fold change| > 0.6. For sequencing of DRH-1(2CARD) animals, differentially expressed genes were determined by adjusted *P* < 0.05 and |log2 fold change| > 1.

Enriched gene categories for significantly upregulated genes were determined using the online tool WormCat 2.0 (http://www.wormcat.com/). Fisher’s exact test was performed in R.

Normalized viral transcript levels were determined using Salmon version 1.11.4[32]. Reads were aligned to the Orsay virus isolate JU1580 genome (NCBI RefSeq assembly = GCF_001402145.1). Transcripts per Million (TPM) values were calculated by running ‘salmon_quant’ on paired-end data.

Differentially expressed genes are listed in S5 to S9 Tables.

### Coimmunoprecipitation

For each sample, 200,000 synchronized L1s were plated onto NGM plates (10 plates, each containing 20,000 worms) and grown at 20°C for 72 h. Animals were fed by top plating *E. coli* OP50-1 as needed to prevent starvation. Adult animals were washed off plates with M9, followed by two additional M9 washes prior to resuspension in 500 µL of ice-cold lysis buffer (50 mM HEPES pH 7.4, 1 mM EGTA, 1 mM MgCl2, 100 mM KCl, 1% glycerol, 0.05% Nonidet P-40, 0.5 mM DTT, and 1x protease inhibitor tablet). Samples were flash frozen by adding worm suspension dropwise into liquid N2.

To prepare protein lysate, frozen worm pellets were ground into a fine powder using a prechilled mortar and pestle. Protein extracts were resuspended in 1-2 ml lysis buffer. To remove debris, extracts were pelleted at 21,000 x g at 4°C for 15 min. The supernatant was filtered using a 0.22 µm PES syringe filter. Protein concentration was measured using a Pierce 660-nm protein assay, and concentrations were adjusted to 1 µg/µL in 1 mL lysis buffer.

For immunoprecipitation, 1 mg of total protein was added to 40 µL of anti-FLAG M2 affinity gel bead suspension (20 µL of packed beads) (Sigma). The lysate/bead mixture was incubated at 4°C overnight with rotation (10 rpm). The bead resin was washed four times with lysis buffer, three times with wash buffer (50 mM HEPES pH 7.4, 1 mM MgCl2, and 100 mM KCl), and three times with 50 mM Tris pH 7.5. The supernatant was removed and beads were store at −80°C.

### LC-MS/MS Analysis

On-bead trypsin digestion was performed to the proteins bound to the beads. Digested peptides were acidified by 0.1% formic acid and analyzed by nanoLC-MS/MS with DIA method.

A Thermo Scientific Vanquish Neo UHPLC system (Buffer A: Water with 0.1% formic acid; Buffer B: 80% acetonitrile with 0.1% formic acid) was used to deliver a flow rate of 400 nL/min to a self-packed capillary C18 (ReproSil-Pur 120 C18-AQ, 1.9um) column (150um ID x 20cm long) with an integrated laser pulled spray tip. Peptides were eluted using a 75min trap-and-elute reverse phase gradient (0–0.1 min 1% B to 7% B, 0.1–61 min 7% B to 35% B, 61–64 min 35% B to 60% B, 64–65 min 60% B to 99% B, 65–69 min 95% B, 69-75 min 1% B).

Mass Spectra were acquired on a Thermo Orbitrap Exploris 480 mass spectrometer operated in positive ion mode with a source temperature of 300°C and spray voltage of 2.1kV. DIA scans were employed with 86 isolation windows of 7 Da (400-1,000 Da), collision energy of 30, and a normalized AGC target of 1000%. The mass resolution was set at 120,000 for MS (profile mode) and 30,000 for MS/MS (centroid mode) scans, respectively. MS scans were performed for every 43 DIA MS/MS scans.

Raw data were analyzed in DIA-NN (version 2.3 Academia)[33, 34]. The spectral library was generated from a UniProt *C. elegans* reference database (UP000001940, 19,824 protein sequences) with common contaminant proteins. The default DIA-NN search parameters were used (Protein inference = “Genes”, Neural network classifier = “Single-pass mode”, Quantification strategy = “QuantUMS (high precision)”, Cross-run normalization = “RT-dependent”, Library Generation = “IDs, RT and IM Profiling” and Speed and RAM usage = “Optimal results”). Mass accuracy and MS1 accuracy were set to 0 for automatic inference. “No share spectra”, “Use isotopologues”, “Heuristic protein inference” and “MBR” were selected. No PTM was selected and 1 tryptic miscleavage is allowed. The false discovery rate (FDR) was set to 1%.

### Proteomics data deposition

The raw spectra of the proteomics data have been deposited in the Mass Spectrometry Interactive Virtual Environment (MassIVE) repository (massive.ucsd.edu/ProteoSAFe/static/massive.jsp, accession ID MSV000102185). FTP download link before publication: ftp://MSV000102185@massive-ftp.ucsd.edu; FTP download link after publication: ftp://massive-ftp.ucsd.edu/v13/MSV000102185/

### Differentially enriched protein (DEP) analysis

The DEP package in R[35] was used to determine differentially expressed proteins in DRH-1(2CARD)::mScarlet samples relative to the mScarlet control. Peptide spectral counts were filtered to remove duplicates and keep only peptides that were identified in all four replicates of at least one condition. Filtered counts were normalized using variance stabilizing normalization. To impute missing values, the MinProb method was used to randomly sample values from a Gaussian distribution centered on a minimal value from the dataset. The fold change ratio and adjusted *P* values were calculated. Significantly enriched proteins were determined based on adjusted *P* < 0.01 and |log2 fold change| > 4. Volcano plots were constructed using the ‘EnhancedVolcano’ package in R. DEP analysis is listed in S4 Table.

### Imaging

All imaging was performed on fixed samples. Images shown in Figs. 1A, 2A, 2C, 4A, and 4B were captured using a Zeiss LSM900 Airyscan confocal microscope and Zen 3.7 software. Images were collected using Airyscan imaging mode SR-4Y and processed using the Airyscan deconvolution algorithm with automatic filter-strength settings. Images shown in Figs. 1C, 1D, and 2D were acquired using a Zeiss Apotome AxioImager compound microscope with Zen 3.9 software and processed using Apotome deconvolution.

### Foldseek analysis of STA-1

The predicted structure of STA-1 was obtained from Alphafold3. Foldseek release 9 was used to determine structural matches. Structures were superimposed in ChimeraX version 1.11.1.

### STAT phylogeny

Signal transducer and activator of transcription (STAT) protein sequences were retrieved from the UniProt/Swiss-Prot database using the search query (protein_name:“signal transducer”) AND (protein_name:“activator of transcription”), filtered to include only entries with a UniProt annotation score of 5[36]. Duplicate sequences were filtered out, and isoforms (same header name, different sequence) were retained and assigned a numerical version suffix (e.g., v2, v3) to ensure unique identifiers across the dataset. A total of 229 final sequences were used for the analysis.

For analysis of full-length STAT proteins, all 229 preprocessed sequences were aligned using MAFFT version 7.525[37] with the parameters (--localpair --maxiterate 1000). The resulting multiple sequence alignment was trimmed using trimAl version 1.5[38] with the automated heuristic trimming mode (-automated1). Phylogenetic inference was performed using IQ-TREE3 version 3.1.1[39] with automatic substitution model selection (-m TEST) and 1,000 bootstrap replicates (-B 1000). The tree was rooted on the *Dictyostelium discoideum* STAT sequence, *dstA*. Phylogenetic trees were visualized in R using the packages ggtree version 4.2.0 and ggtreeExtra version 1.22. Trees were displayed as circular cladograms (branch.length = “none”).

### Statistics

All statistical analyses were performed in R. Q-Q plots were used to assess normality of the data, and parametric tests were used when appropriate. A nonparametric test was applied to data that did not meet the assumptions for a parametric test.

## Acknowledgements

We thank Pooranachithra Murugesan and Andrew Chisholm for support with confocal imaging. We thank the UCSD Goeddel Family Technology Sandbox for providing the Thermo Scientific Vanquish Neo UHPLC system and Exploris 480 mass spectrometer used in this research to generate the proteomics data, as well as Steve Briggs for his support. We thank Marisa Tsunoda, Eilish Murphy and Joshua Joseph for generating *jyIs41*. This publication includes data generated at the UC San Diego IGM Genomics Center utilizing an Illumina NovaSeq X Plus that was purchased with funding from a National Institutes of Health SIG grant (#S10 OD026929). Some strains used in this study were provided by the Caenorhabditis Genetics Center (CGC), which is funded by NIH Office of Research Infrastructure Programs (P40 OD010440). This work was supported by NIH under R01 AG052622, GM114139, and AI176639 to E.R.T., by 5F31AI176729 to L.E.B., and by NIGMS/NIH award K12GM068524 to T.D.B.

## Supporting Information

**Fig. S1.**
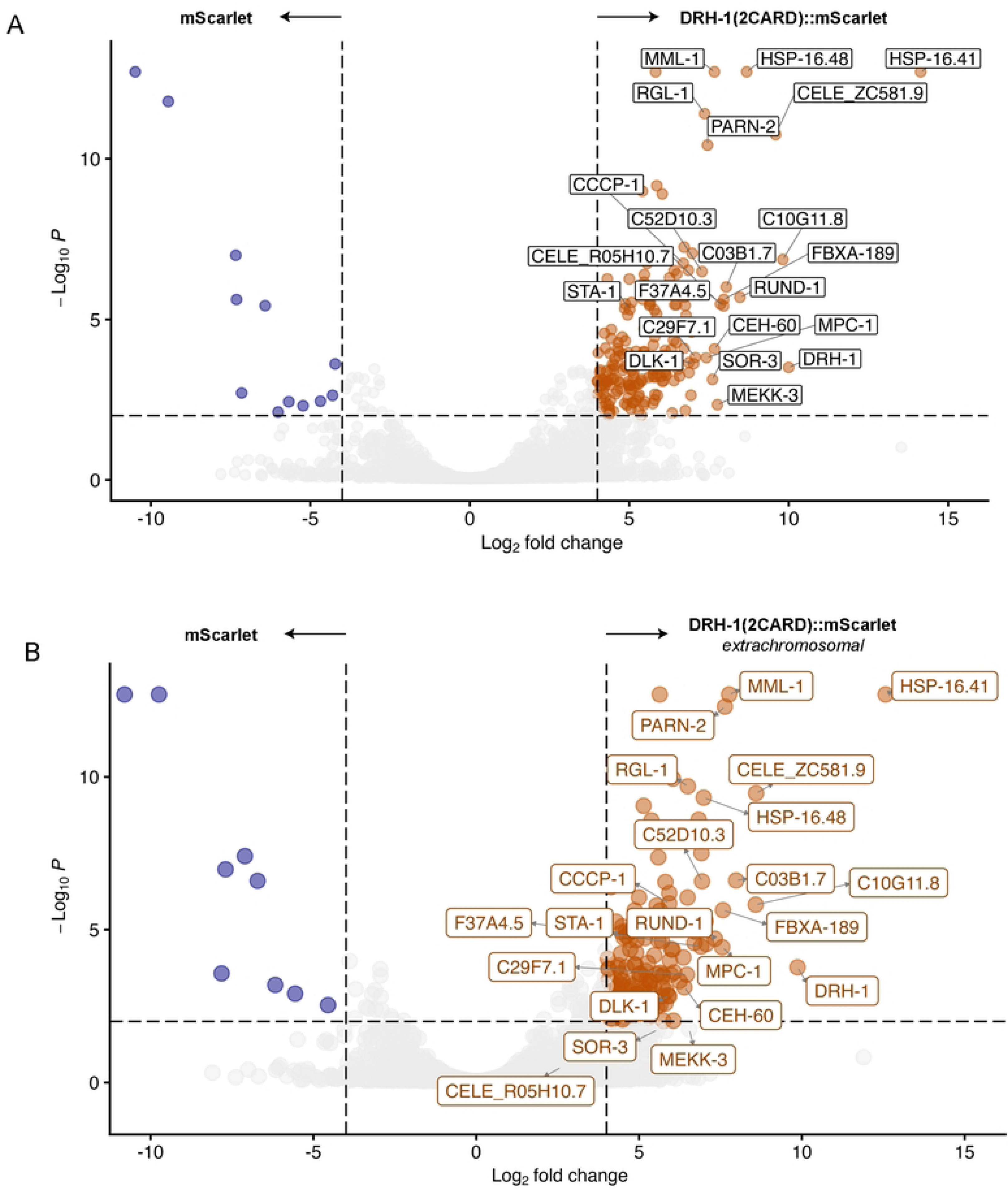
DRH-1(2CARD) protein interactors identified by co-IP/MS. (A) Differentially enriched proteins when comparing integrated DRH-1(2CARD) (*jyIs37) vs.* mScarlet control (*jyIs41).* Both constructs expressed in *rde-1(ne300)* mutant background. (B) Differentially enriched proteins when comparing extrachromosomal DRH-1(2CARD) (*jyEx305) vs.* mScarlet control (*jyIs41).* Extrachromosomal DRH-1(2CARD) is expressed in a WT background. Differentially enriched proteins determined by |log2FC| > 4 and p < 0.01.

**Fig. S2.**
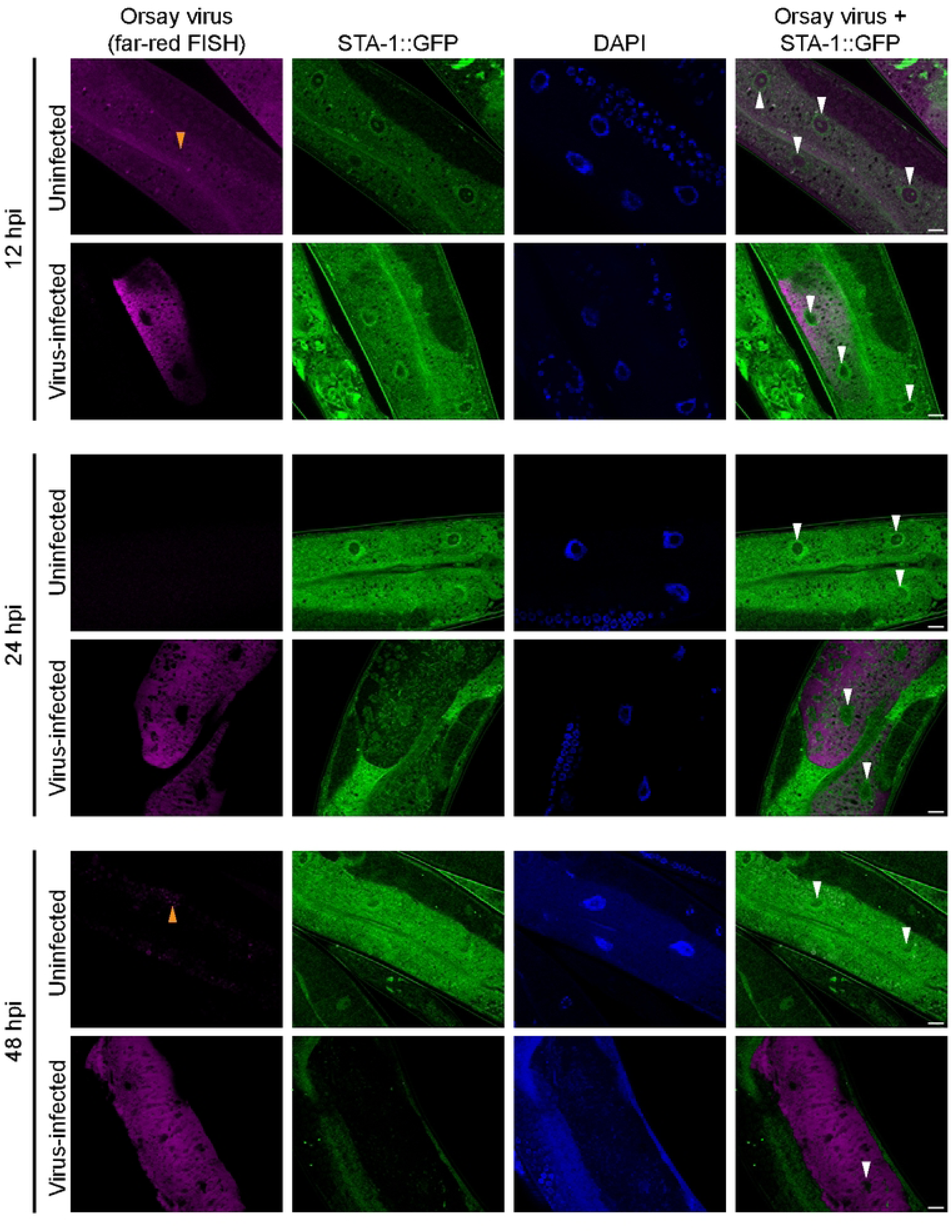
Time course of changes in STA-1::GFP expression upon viral infection. Representative images of uninfected or virus-infected adult animals at 12, 24, and 48 hpi. Animals were infected at the L4 stage. Viral infection was visualized using Quasar 670-conjugated (far-red) FISH probes targeting the Orsay virus genome. Nuclei were counterstained using DAPI (blue). White arrowheads indicate nuclei; orange arrowheads indicate autofluorescence. Scale bar = 10 µm.

**Fig. S3.**
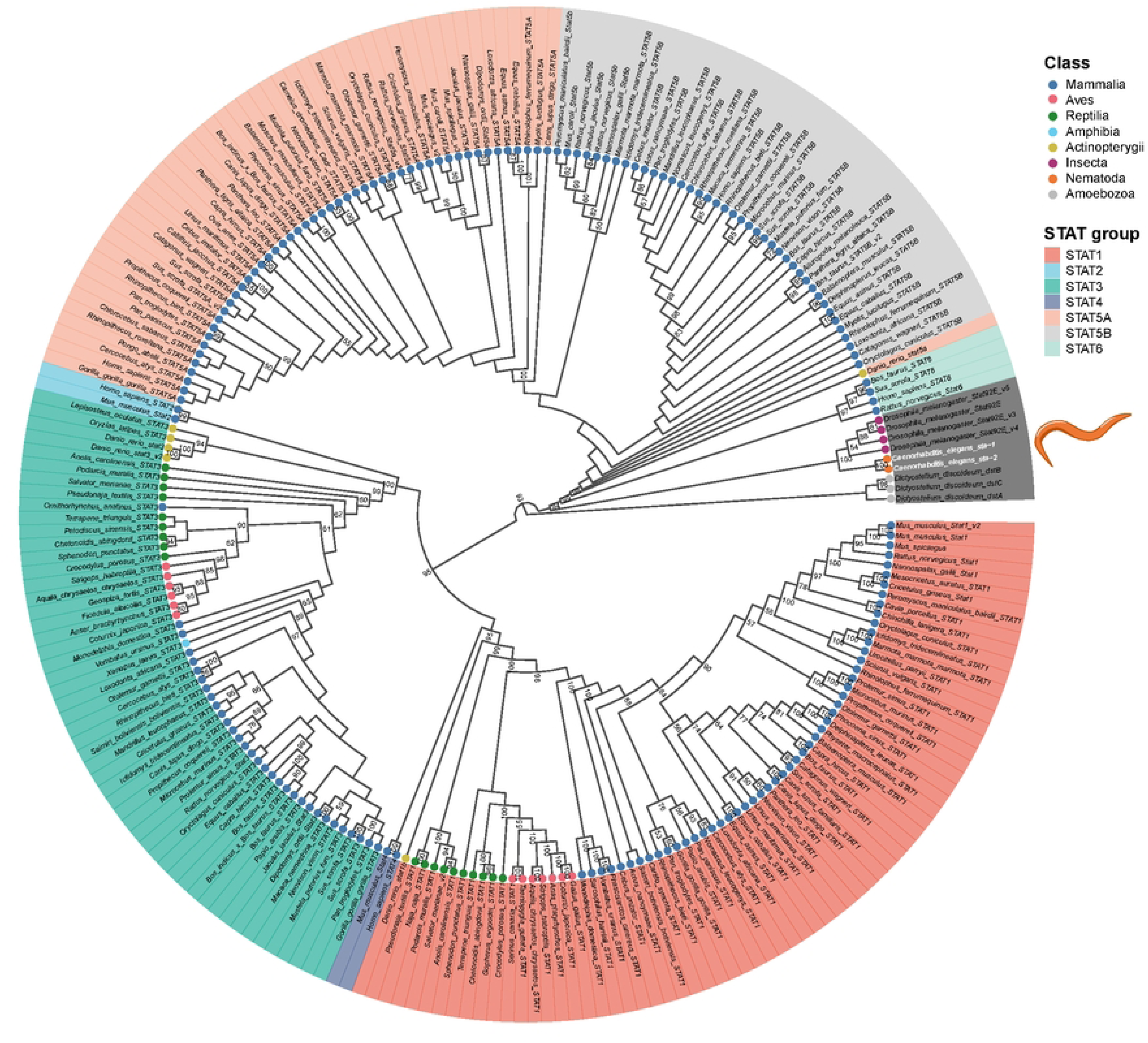
STAT phylogenetic analysis. Phylogenetic tree constructed with amino acid sequences of full-length STAT proteins. Text shading indicates mammalian STATs and annotated homologs. Node coloring indicates taxonomic rank. Bootstrap values over 50 are listed at nodes.

**Fig. S4.**
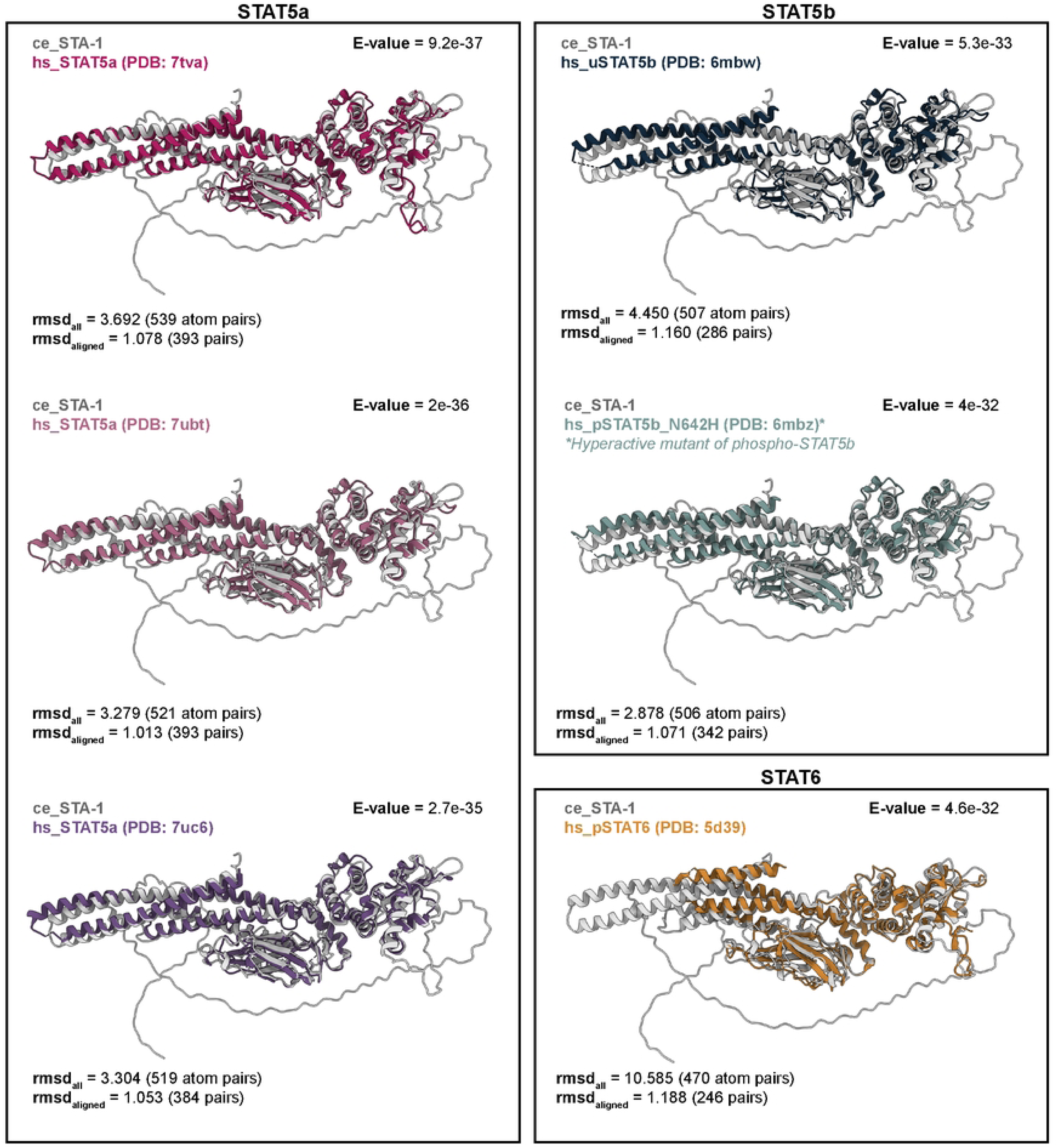
Foldseek identifies human STAT5a/STAT5b as top structural matches for STA-1 Alphafold predicted structure. Overlay of STA-1 Alphafold structure (white) and top 6 structural matches: STAT5a (pink/purple), STAT5b structures (blue/green), and STAT6 (yellow). Lower E-values indicate higher similarity between structures. Root mean square deviation (RMSD) is reported for all atoms in both structures (RMSD_all_) or for atoms at aligned amino acid residues (RMSDaligned).

**Fig. S5.**
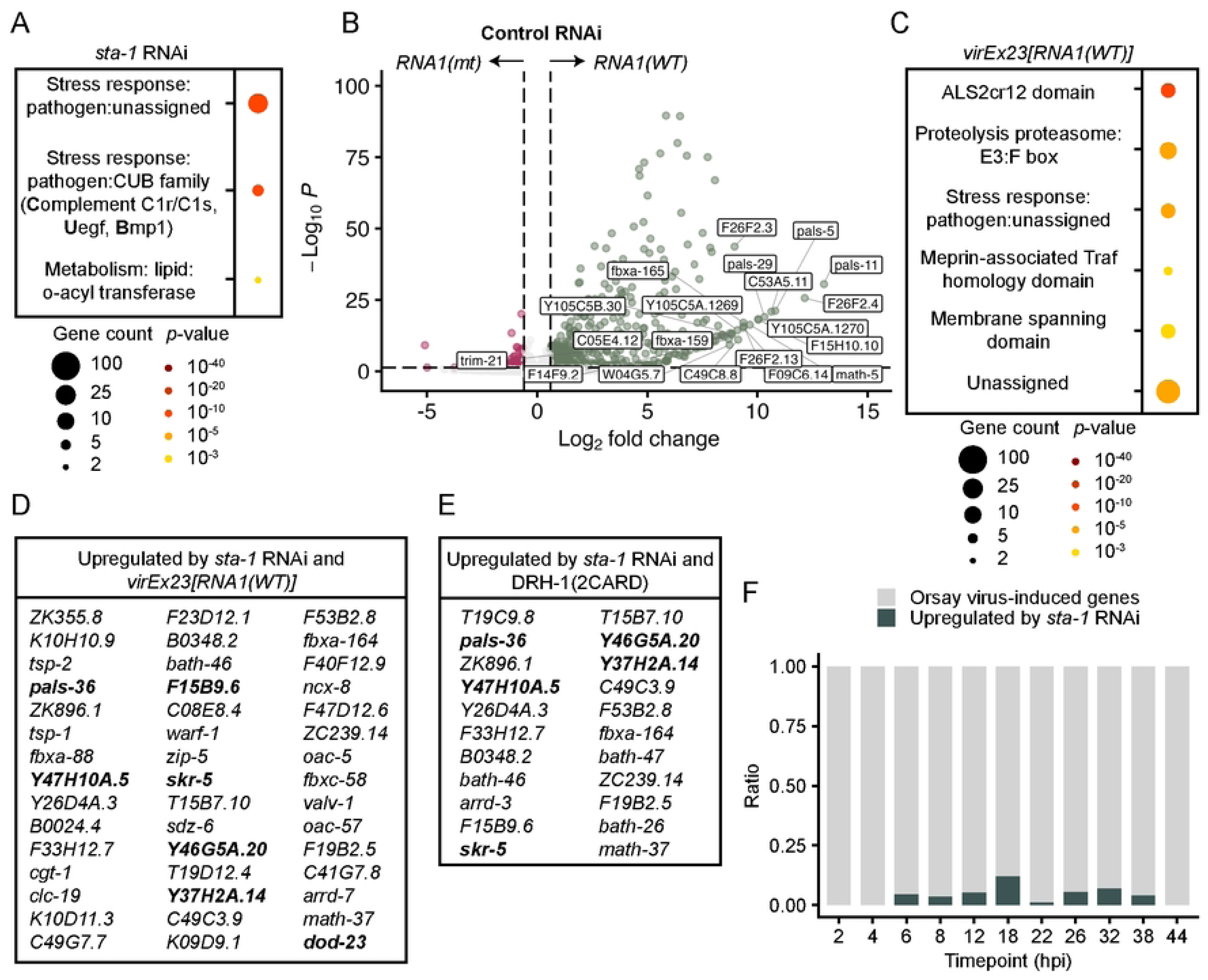
STA-1 negatively regulates genes induced by viral RNA1 replicon and infection. (A) WormCat analysis of upregulated genes in *sta-1* vs. control RNAi in *virEx26[RNA1(mt)]* background. (B) Volcano plot of differentially expressed genes (DEGs) in *virEx23[RNA1(WT)]* vs. *virEx26[RNA1(mt)]*. Both groups were treated with control RNAi. Upregulated (green) and downregulated (magenta) genes determined by |log2FC| > 0.6 and p < 0.05. Labels indicate top 20 upregulated genes. (C) WormCat analysis of upregulated genes in *virEx23[RNA1(WT)]* vs. *virEx26[RNA1(mt)]* animals treated with control RNAi. For (A) and (C) gene counts and p-values are displayed in scaled bubble charts. Enrichment scores were determined using a false discovery rate cutoff of 0.01. (D) Genes upregulated by both *sta-1* RNAi and *virEx23[RNA1(WT)].* (E) Genes upregulated by both *sta-1* RNAi and DRH-1(2CARD). (F) The proportion of STA-1-regulated genes (green) ranges from 0 to 0.14 of total upregulated virus response genes at a given timepoint during the infection time course (Castiglioni et al. 2024).

### Supplementary Tables

**S1 Table.** *C. elegans* strains used in this study

**S2 Table.** DNA constructs used in this study

**S3 Table.** Primers used in this study

**S4 Table.** Co-immunoprecipitation mass spectrometry results

**S5 Table.** Differentially expressed genes in *virEx23* L4440 vs *virEx26* L4440

**S6 Table.** Differentially expressed genes in *virEx23 sta-1* RNAi vs *virEx23* L4440

**S7 Table.** Differentially expressed genes in *virEx23 sta-1* RNAi vs *virEx26 sta-1* RNAi

**S8 Table.** Differentially expressed genes in *virEx26 sta-1* RNAi vs *virEx26* L4440

**S9 Table.** Differentially expressed genes in *jyIs37* vs *jyIs41*

